# Global Biobank Engine: enabling genotype-phenotype browsing for biobank summary statistics

**DOI:** 10.1101/304188

**Authors:** Gregory McInnes, Yosuke Tanigawa, Chris DeBoever, Adam Lavertu, Julia Eve Olivieri, Matthew Aguirre, Manuel Rivas

## Abstract

Large biobanks linking phenotype to genotype have led to an explosion of genetic association studies across a wide range of phenotypes. Sharing the knowledge generated by these resources with the scientific community remains a challenge due to patient privacy and the vast amount of data. Here we present Global Biobank Engine (GBE), a web-based tool that enables the exploration of the relationship between genotype and phenotype in large biobank cohorts, such as the UK Biobank. GBE supports browsing for results from genome-wide association studies, phenome-wide association studies, gene-based tests, and genetic correlation between phenotypes. We envision GBE as a platform that facilitates the dissemination of summary statistics from biobanks to the scientific and clinical communities. GBE currently hosts data from the UK Biobank and can be found freely available at *biobankengine.stanford.edu*.

## 1. Introduction

Population-scale biobanks linking rich phenotype and molecular data are transforming the landscape of biomedical research. UK Biobank, a long-term prospective cohort study, has collected array-genotyped variants from 500,000 individuals and linked it with medical records, activity monitors, imaging, and survey data (Sudlow et al., 2015). Availability of these data enables researchers to perform analyses across a broad range of phenotypes at an unprecedented scale (Bycroft et al., 2017).

The value of large sequencing and genotyping efforts lies not only in primary publications but also in the dissemination of summary statistic data to the scientific community. Other large-scale efforts to sequence and analyze genetic data, such as ExAC and gnomAD (Lek et al., 2016), have made data available to the scientific community at large available via web browsers (Karczewski et al., 2017). Browsers serve as an effective communication tool that enable researchers around the world to interrogate genetic statistics of interest. Often, these tools limit the information shared to summary statistics which confers a decreased privacy risk for individuals included in the study (Erlich & Narayanan, 2014) and limits the computational resources required to interrogate the data. However, to date no such tool exists that offers researchers the opportunity to study the relationship between genotype and phenotype.

Here we present Global Biobank Engine (GBE), a web-based tool that presents summary statistics resulting from analysis of genotype-phenotype relationships derived from data in population-scale biobanks. This tool serves as a means to communicate scientific discoveries to the research and clinical communities without requiring sharing of individual-level data. In particular, we present results from genome-wide association studies (GWAS) and phenome-wide association studies (PheWAS) for White British individuals (n = 337,199) in UK Biobank, gene-level phenotype associations (including rare variant aggregate analysis), genetic correlations, and others. Results for each analysis are precomputed allowing for rapid browsing. Phenotypes currently available in the browser are those made available by UK Biobank, including cancer, disease status, family history of disease, medication, quantitative measures, as well as computational grouping of phenotypes based on self-reported data and ICD 10 codes from hospital in-patient record data (as described in (DeBoever et al., 2017)).

### 2. Features

Global Biobank Engine serves as a platform to host summary statistics that explore different facets of biobank data. Here we highlight the methods available in GBE.

### 2.1. Phenotype page

The phenotype page presents a summary of the results of a GWAS run for a phenotype of interest. The first part of the page displays relevant data such as the sample count included in the GWAS as well as links to other analyses related to this phenotype (Fig. 1.A1). Next, the Manhattan plot is displayed including all variants with p-value < 0.001 (Fig. 1.A2). Finally, a table is included with detailed information for each variant is included. The table can be subsetted by all variants, protein truncating variants (PTVs) only, or both PTVs and missense variants.

**Fig. 1.**
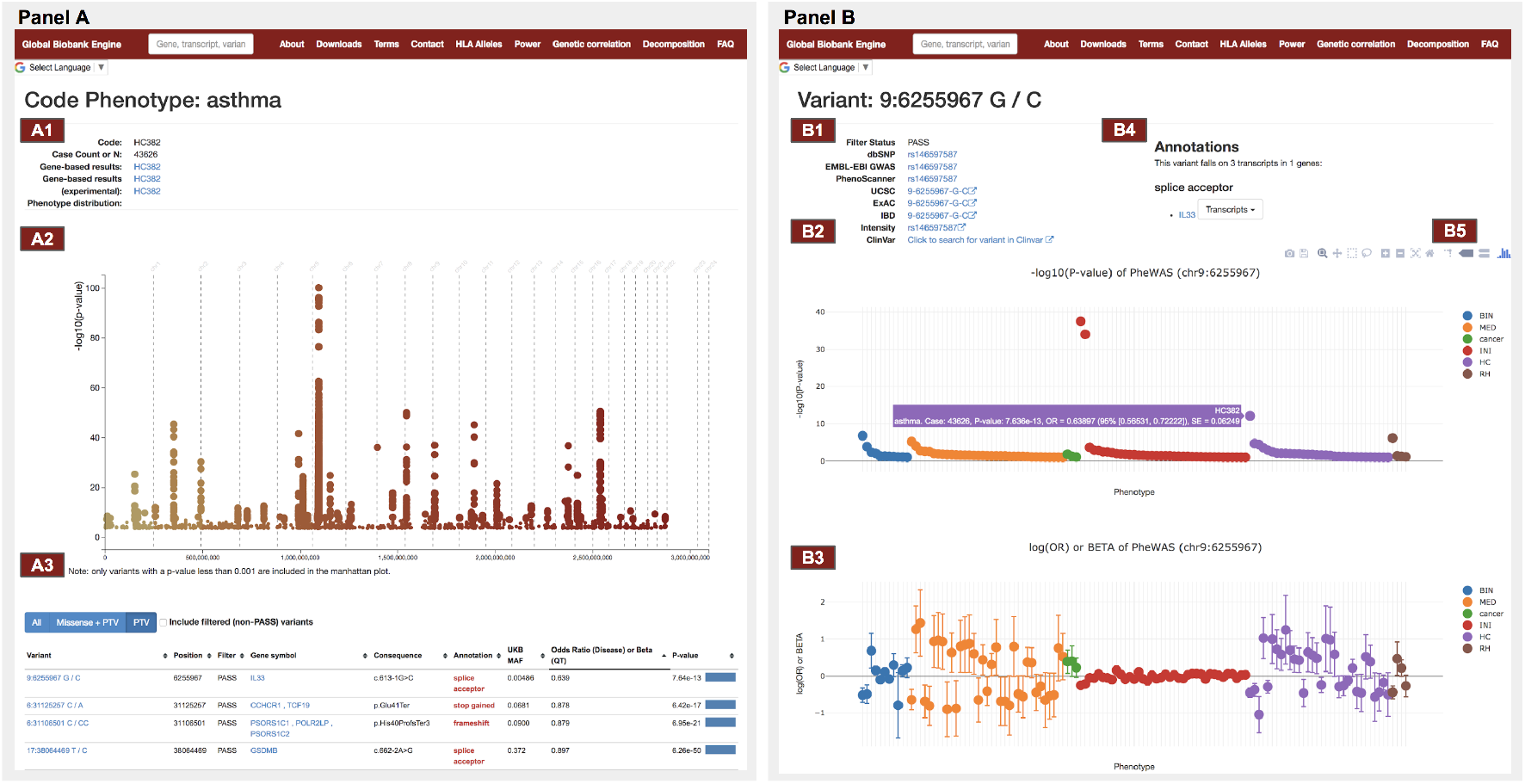
GBE screenshots of phenotype page (left) and variant page (right). Shown here is the phenotype page for asthma in the UK Biobank and the variant page for the protein-truncating variant rs146597587 in *IL33* found to protect against asthma. (A1) Summary of phenotype information including sample count and links to other phenotype-specific analyses. (A2) Manhattan plot displaying significance of association of each genotyped variant with a p-value < 0.001. (A3) Detailed variant information is summarized in a table with links to variant pages. (B1) Variant summary and link-outs to external references. (B2) Manhattan plot for a PheWAS of rs146597587. Phenotypes are binned by category. (B3) Effect size estimate plot of the log(OR) or beta values for rs146597587 for each phenotype. (B4) Variant annotations and links to associated genes. (B5) Figures can be saved locally and manipulated using the set of tools provided.

### 2.2. Variant page

The variant page presents the annotation of a genetic variant (Fig. 1.B4), links to external resources (Fig. 1.B1), and two plots summarizing the results from PheWAS analysis of the variant. The PheWAS Manhattan plot on the top presents the statistical significance of associations (Fig. 1.B2) while the effect size plot on the bottom presents the log odds-ratio and regression coefficient for binary and continuous traits, respectively (Fig. 1.B3). The phenotypes in the plots are sorted by their category and can be subset by p-values. One can export the plots to image files to facilitate scientific communication (Fig. 1.B5).

### 2.3. Gene page

The gene page presents a summary of all genotype-phenotype statistics related to a single gene. This page includes a Manhattan plot which displays each variant in the gene region and the phenotype with the lowest p-value for that variant as well as a table summarizing additional variant information. The page also includes a figure showing the top 5 most related phenotypes by rare variant aggregate analysis with a link to a full description of the rare variant aggregate statistics for that gene.

### 2.4. Genetic correlation page

GBE includes an interactive application for browsing genetic correlation estimates for pairs of traits from the UK Biobank. Genetic correlations have been estimated by applying the multivariate polygenic mixture model (MVPMM) to GWAS summary statistics for more than one million pairs of traits and can be visualized using the app (DeBoever et al., 2018). Users can filter the phenotypes and results that are displayed by the app. MVPMM also estimates other genetic parameters including polygenicity and scale of effects which can be seen by mousing over the plot.

### 2.5. HLA alleles

The HLA alleles page shows posterior probabilities of causal associations between 175 HLA allelotypes and 270 diseases in the UK Biobank. For each allelotype there is a plot showing the log odds ratio with a 95% confidence interval for each associated phenotype with posterior probability greater than 0.7. Users can also view donut charts displaying the frequencies of allelotypes at each locus.

## 3. Implementation

GBE extends the ExAC browser (Karczewski et al., 2017) which is built in Python and utilizes Flask framework and d3 and plot.ly for plot rendering. One key change made in our implementation is the use of a SciDB backend to host the summary statistic data presented in the browser (Rivers, 2017). We found SciDB to have superior performance with the large amount of data that needs to be stored and queried. GBE is hosted within a Docker virtual environment on Google Cloud.

## 4. Availability

GBE browsing capabilities are now publicly available at biobankengine.stanford.edu.

## 5. Acknowledgements

This research has been conducted using the UK Biobank Resource under Application Number 24983. We thank all the participants in the UK Biobank study. MAR is supported by Stanford University and a National Institute of Health center for Multi - and Transethnic Mapping of Mendelian and Complex Diseases grant (U01 HG009080), National Human Genome Research Institute GSP Coordinating Center (U24 HG008956) and the MoTr-PAC Bioinformatics Center (1U24EB023674-01). MAR is a Faculty Fellow at the Stanford Center for Population Health Sciences. G.M. and A.L. are supported by BD2K grant number T32 LM 012409. C.D. is supported by a postdoctoral fellowship from the Stanford Center for Computational, Evolutionary, and Human Genomics and a seed grant from the Stanford ChEM-H Institute. Y.T. is supported by Funai Overseas Scholarship from Funai Foundation for Information Technology and the Stanford University Biomedical Informatics Training Program. The primary and processed data used to generate the analyses presented here are available in the UK Biobank access management system (https://amsportal.ukbiobank.ac.uk/) for application 24983, ‘‘Generating effective therapeutic hypotheses from genomic and hospital linkage data” (http://www.ukbiobank.ac.uk/wp- content/uploads/2017/06/24983-Dr-Manuel-Rivas.pdf), and the results are displayed in the Global Biobank Engine (https://biobankengine.stanford.edu). We would also like to thank Christopher Chang for his work optimizing PLINK2.

